# Silencing of the CSNK2β gene by siRNA inhibits invasiveness and growth of MDA-MB-231 cells

**DOI:** 10.1101/311092

**Authors:** Shibendra Kumar Lal Karna, Bilal Ahmad Lone, Faiz Ahmad, Nerina Shahi, Yuba Raj Pokharel

**Author notes:** Corresponding author: Yuba Raj Pokharel.

## Abstract

**Background:** Breast cancer is most common cancer and accounts for one-fourth of all cancer diagnoses worldwide. Treatment of triple-negative breast cancer is major challenge and identification of specific drivers is required for targeted therapies. The aim of our present study is to elucidate the therapeutic potential of CSNK2β silencing in triple negative breast cancer MDA-MB-231 cell.

**Methods:** CSNK2ß gene has been knockdown using siRNA and silencing was estimated by both real time and western blot. Cell Titer-Glo (CTG) and colony formation assay and wound healing assay, cell cycle analysis by flow cytometry was performed to assess the role of CSNK2ß in cell proliferation, migration, cell cycle, and oncogenesis. Morphological assessment of nuclear condensation, apoptosis by Hoechst staining and measurement of intracellular ROS production was examined using fluorescence microscopy. Real time PCR and western blot was done to study the expression of genes related to cell proliferation, survival, metastasis, apoptosis, and autophagy.

**Results:** Silencing of CSNK2ß in MDA-MB-231 cells resulted in decreased cell viability, colony formation, and migratory potential. Cell cycle analysis showed that growth inhibitory effect was mediated by arresting the cells in G2/M phase. Furthermore, we demonstrated that silencing of CSNK2β increased the nuclear condensation and intracellular ROS production. CSNK2β regulates the expression of BAX, Bcl-xL, caspase 3, Beclin-1, LC3-I, p-ERK, p38-α, c-Myc, MAPK, c-Jun, NF-ĸB, β-catenin, E2F1, PCNA. We have also shown the functional relationship between CSNK2β, PIN1, and PTOV1 by western blotting. We have first time reported that silencing CSNK2β using siRNA can inhibit invasiveness and proliferation of MDA-MB-213 cells.

**Conclusion:** Our results suggested that CSNK2β silencing may offer future therapeutic target in triple negative breast cancer.

## Background

Casein Kinase 2(CSNK2), a highly conserved, multifunctional serine/threonine protein kinase, is critically important for the regulation of different processes in eukaryotes, such as proliferation, differentiation, and apoptosis [1–3]. CSNK2 is ubiquitously expressed in all tissues but its level is elevated in tumor tissues including prostate [4], mammary gland [5], head and neck [6], lungs [7] and kidney [8]. CSNK2 possess a heterotetrameric conformation with two catalytic and two regulatory subunits [9]. CSNK2β supports the structure of the tetrameric complex, augments catalytic activity and stability of CSNK2 and can also function dependently with other catalytic subunits [10–13, 9].

In a mammalian cell, CSNK2β is phosphorylated at Ser 209 at its autophosphorylation site and at Ser 53B, in a cell cycle-dependent manner and *in vitro* by p34^cdc5^ [14, 15]. CSNK2ß is responsible for recruitment of CSNK2 substrates or potential regulators such as Nopp140, p53, Fas-associated factor-1 (FAF-1), topoisomerase II, CD5 and fibroblast growth factor-2 (FGF-2) [16–22]. Yeast two-hybrid interaction showed that CSNK2β interact with c-Mos and A-Raf protein kinases [23–25]. Ectopic expression of CSNK2β in mouse 3T3-L1 adipocytes and in CHO cells increased proliferation [26]. The proliferative effects of CSNK2β vary in different cell lines. Deletion of gene encoding CSNK2ß in mice leads to a failure in development [27].

To understand its physiological importance in the regulation of multiple candidate target proteins, we focused our study on the role of CSNK2ß in the tumorigenesis of human breast cancer (MDA-MB-231 cell) in vitro. In the present study, we used the RNA interference strategy to knockdown the CSNK2ß gene and study the gross oncogenic activity in *an in-vitro* cell-based system. We evaluated its clonogenic, invasive, proliferative and apoptotic properties in MDA-MB-231 cells using siRNA. We found that CSNK2β regulates the cell proliferation by targeting NF-ĸB, Wnt, JNK and MAPK pathway proteins and also modulates the expression of PIN1 and PTOV1 oncogene. Our findings suggest that CSNK2β can be used as a novel target for breast cancer therapy.

## Materials and methods

### Reagents

Lipofectamine RNAiMAX, TRIzol, Propidium Iodide, RNase were purchased from Invitrogen Corp (Carlsbad, CA, USA). siRNA was obtained from Qiagen (Hilden, Germany). Cell culture reagents and flasks were purchased from HiMedia (France) and Corning Inc (Corning, New York, USA). SYBR Green was purchased from Bio-Rad (Hercules, California). Antibodies were obtained from Santa Cruz Biotechnology (Dallas, Texas, USA), Cloud-Clone Corp. (Houston, USA). Cell Titer-Glo reagent was purchased from Promega Corp (Madison, Wisconsin, USA).

### Cell culture

MDA-MB-231 cell was purchased from National Centre for Cell Science (Pune, India). The cells were cultured in L-15 medium supplemented with 10% FBS, penicillin (100 unit/ml) and streptomycin (100μg/ml). The cell culture was maintained at 37°C in humidified air containing 5% CO_2_.

### Transfection

Cells were cultured in 6 well plates one day before siRNA transfection. We used 25 nanomolar of each siRNA and made complex in Opti-MEM media. Similarly, the complex of Lipofectamine RNAiMAX (4 μl/each well) and Opti-MEM was made and incubated for 5 minutes at room temperature. After that, both the complexes were mixed in 1:1 proportion and incubated for 25 minutes at room temperature. Cells were treated with Opti-MEM-siRNA Lipofectamine complex and incubated at 37°C for 72 hours. The sequences used in siRNA and the target sequence for the genes in our study are mentioned below.

Scramble siRNA (Negative control)

Target sequence 5’-AATTCTCCGAACGTGTCACGT-3’

Sense strand 5’-UUCUCCGAACGUGUCACGUdT dT-3’

Antisense strand 5’-ACGUGACACGUUCGGAGAAdT dT-3’

CSNK2β siRNA

Target sequence 5’-CAGGACAAATTTAATCTTACT-3’

Sense strand 5’-GGACAAAUUUAAUCUUACUTT-3’

Antisense strand 5’-AGUAAGAUUAAAUUUGUCCTG-3’

Pin1 siRNA

Target sequence 5’-GACCGCCAGATTCTCCTTAA-3’

Sense strand 5’-CCGCCAGAUUCUCCCUUAATT-3’

Antisense strand 5’-UUAAGGGAGAAUCUGGCGGTC-3’

PTOV1 siRNA

Target sequence 5’-CCGGCGTGTCATTGCCAACCA-3’

Sense strand 5’-GGCGUGUCAUUGCCAACCATT-3’

Antisense strand 5’-UGGUUGGCAAUGACACGCCGG-3’

### Construction of CSNK2ß Overexpression plasmid and transfection

CSNK2β was cloned into pcDNA 3.1(+) expression vector. 2.5 μg of empty vector and Overexpression clone was used for transfection. Briefly, 1:1 complex of lipofectamine 2000 plus Opti-Mem and plasmid plus Opti-Mem were made and incubated for 20 minutes. The cell plating number was similar to siRNA transfection.

### Cell viability assay

Cell viability was accessed with CTG assay (Promega, Madison, WI). Briefly, MDA-MB-231 cells were seeded in 96 well white culture plate at a density of 1000 cells per 180 μl of medium per well with 20 μl of siRNA complexes for CSNK2β and incubated at 37°C, 5% CO_2_ and incubated for 24 hours. On the next day, the media containing the complex was changed with the fresh media and further incubated till 96 hours. The cells were treated in quadruplets with respective siRNA. The reagents were prepared according to manufacturer’s protocol. After incubation 100 μl of fresh media was added to each well followed by 100 μl of reagent and kept on a shaker for 2 minutes to induce the cell lysis. The plate was incubated for 10 minutes at room temperature to stabilize the luminescence signal. Luminescence was measured using microplate ELISA reader (Bio Tek, Winooski, Vermont, US).

## COLONY FORMATION ASSAY

MDA-MB-231 cells were transfected with Scramble and CSNK2β siRNA and incubated for 48 hours. After the cells were trypsinized, collected and counted. 1000 cells /pell were taken from each Scramble and Csnk2β transfected samples and seeded in 6 well plate. The cells were allowed to grow for 3 weeks until 80-90% confluence was observed. Then the cells were washed with DPBS, fixed with 3.7% formaldehyde for 30 min and stained with 0.4% crystal violet. The cells washed DPBS for 2-3 times and allowed to dry. The colonies were counted using Image J software.

### Wound healing assay

MDA-MB-231 cells were plated in 6 well plates (4×10^5^ cells/well) and transfected with Scramble and CSNK2β siRNA as mentioned above. After the cells reach 90% confluence, a wound was made in the center of the plate by scrapping the cells monolayer with 10μl tip. The cells were washed with sterile DPBS to remove the detached cells. The width of wound area was measured at different time points using an inverted microscope. Densitometry analysis of wound area was carried by using Image J software

### Cell cycle analysis

2.5□10^5^ cells were seeded in 6 well plates and transfection was done as mentioned before. The cell cycle analysis was performed 48 hours post-transfection using BD FACS Verse ™. The cells were pooled out and fixed with ice-cold 70% ethanol and kept overnight at 4°C. On the next day, the cells were washed with DPBS for two times. The cells were further digested with RNase A (50 U/ml) for 1 hour and then stained with PI (20 μg/ml). The samples were incubated at 4°C for 30 min and analyzed by flow cytometer.

### Western blotting

MDA-MB-231 cells were cultured in 6 well plates. On the next day, after the cells reached the confluence of 70-80%, transfected with Scramble and CSNK2β siRNA and incubated for 72 hours. After incubation, the cells were washed twice with ice-cold DPBS and then lysed with sodium dodecyl phosphate (SDS) lysis buffer. The samples were heated for 5 minutes at 100°c. The samples were run on SDS-PAGE, transferred to PVDF membrane (100 volts for 1 hour). The membranes were blocked using 5% skim milk in Tris buffer saline with 0.1% tween20 (TBST) for 2 hours at room temperature. The blots were incubated with primary antibodies overnight at 4°c on a rocker followed by incubation with HRP conjugated secondary antibodies. The blots were developed using enhanced chemiluminescence (ECL, Biorad). β actin was developed as a loading control.

### RNA extraction, cDNA preparation, and real time PCR

Cells were transfected for 72 hours with the respective siRNAs and RNA was extracted using TRIzol lysis reagent using manufacturers protocol. 2μg of RNA was taken and pretreated with DNase followed by cDNA synthesis with oligoDT primers. The thermocycler condition was 65° c, 5 min, 42°c, 60 min and 72°c (final extension). Real time PCR was done with iTaq SYBR green mix (Biorad) using ABI 700,(Invitrogen). PCR was carried out at following conditions.

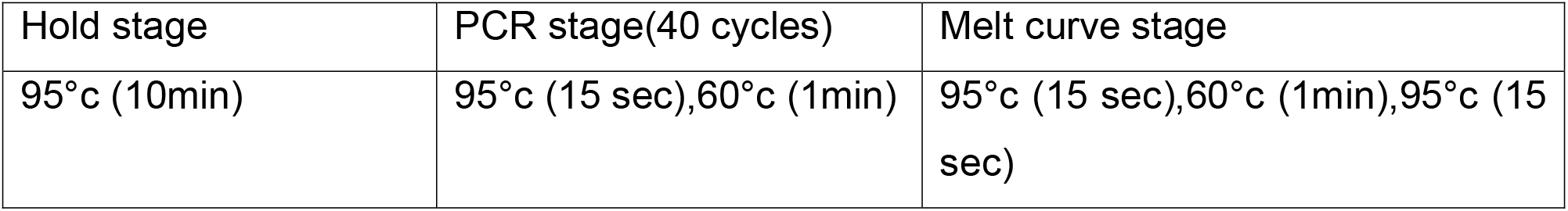

**Table 1.**
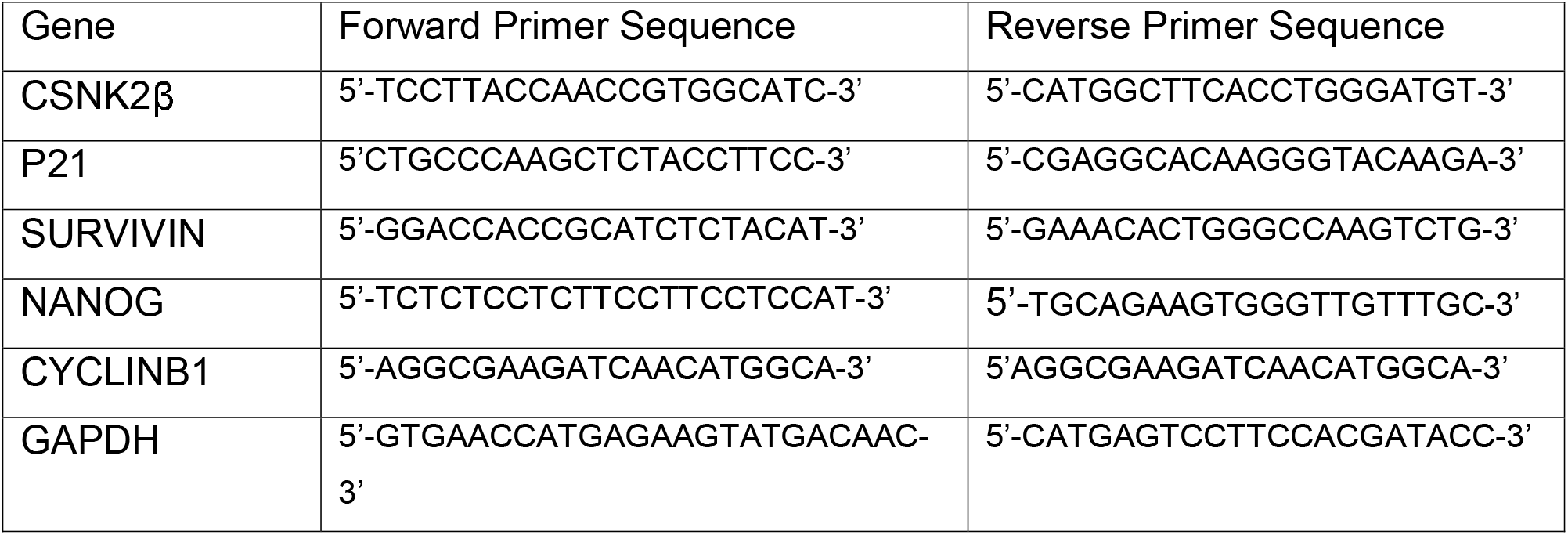
List of sequence of primers for Real time PCR

### Hoechst staining

1.5□10^5^ cells were seeded in 6 well plates. On the next day, the cells were transfected with Scramble and CSNK2β siRNA and incubated for 72 hours. Then the cells were stained with Hoechst 33342 (2μg) stain for 15 minutes in dark. Cells were once washed with DPBS and the images were taken using Nikon fluorescent microscope.

### Measurement of the intracellular ROS production

Intracellular ROS generation was accessed by fluoroprobe DCF-DA (Invitrogen). Approximately 1.5□10^5^ cells were cultured in 6 well plates and transfected with the Scramble and CSNK2β siRNA for 72 hours. The cells were treated with 10 μM CM-H 2 DCFDA for 30 min in the dark at 37°C. Cells were then stained with Hoechst 33342 for 15 minutes in dark at room temperature for nuclear staining. The wells were once washed with DPBS and the images were captured using a fluorescence microscope (Nikon Ti).

### Statistical Analysis

Data were represented as a mean ± standard deviation. The level of significance between two groups was calculated by t-test. P < 0.05 was considered statistically significant.

## Results

### Expression of CSNK2ß is altered in a multitude of tumor types

cBioPortal data shows that CSNK2β has alterations (mutation, amplification, and deep deletion) across the human cancer types. The histogram shows the potential cancer relevance of CSNK2β and high frequency of alteration in Breast (BCCRC xenograft) that prompted us to validate its function in breast cancer (Figure 1) [28].

**Figure 1.**
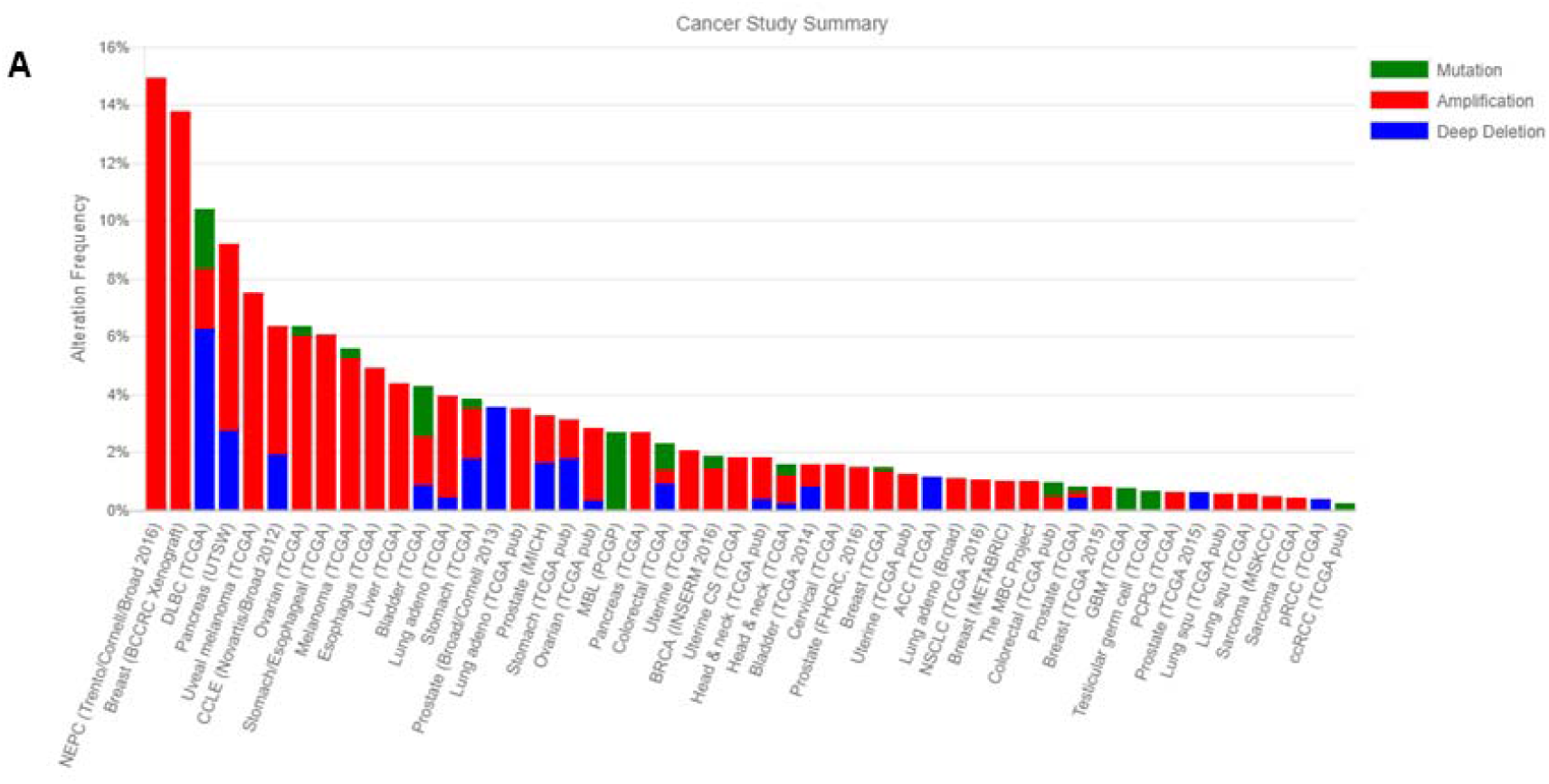
Mutational status of CSNK2β across human cancers based on cBioPortal data. Histogram representing the mutational landscape (mutation, amplification, and deep deletion) of CSNK2ß in different types of cancer-based on cBioPortal data.

### Silencing of CSNK2ß inhibits the proliferation of MDA-MB-231 cells

CSNK2ß gene was silenced by transfecting siRNA in MDA-MB-231breast cancer cell. Silencing of CSNK2β caused the decrease in the expression of the CSNK2β protein (Figure 2A). Cell viability was measured after 96 hours of post-transfection by using CTG reagent which detects the level of adenosine triphosphate (ATP) present in the samples. The growth rate and the viability of CSNK2β silenced cells were significantly lower as compared to scrambled siRNA (p< 0.001) (Figure 2B). These results suggested that CSNK2β play a crucial role in cell proliferation of MDA-MB-231 cells as shown in figure 2B.

**Figure 2.**
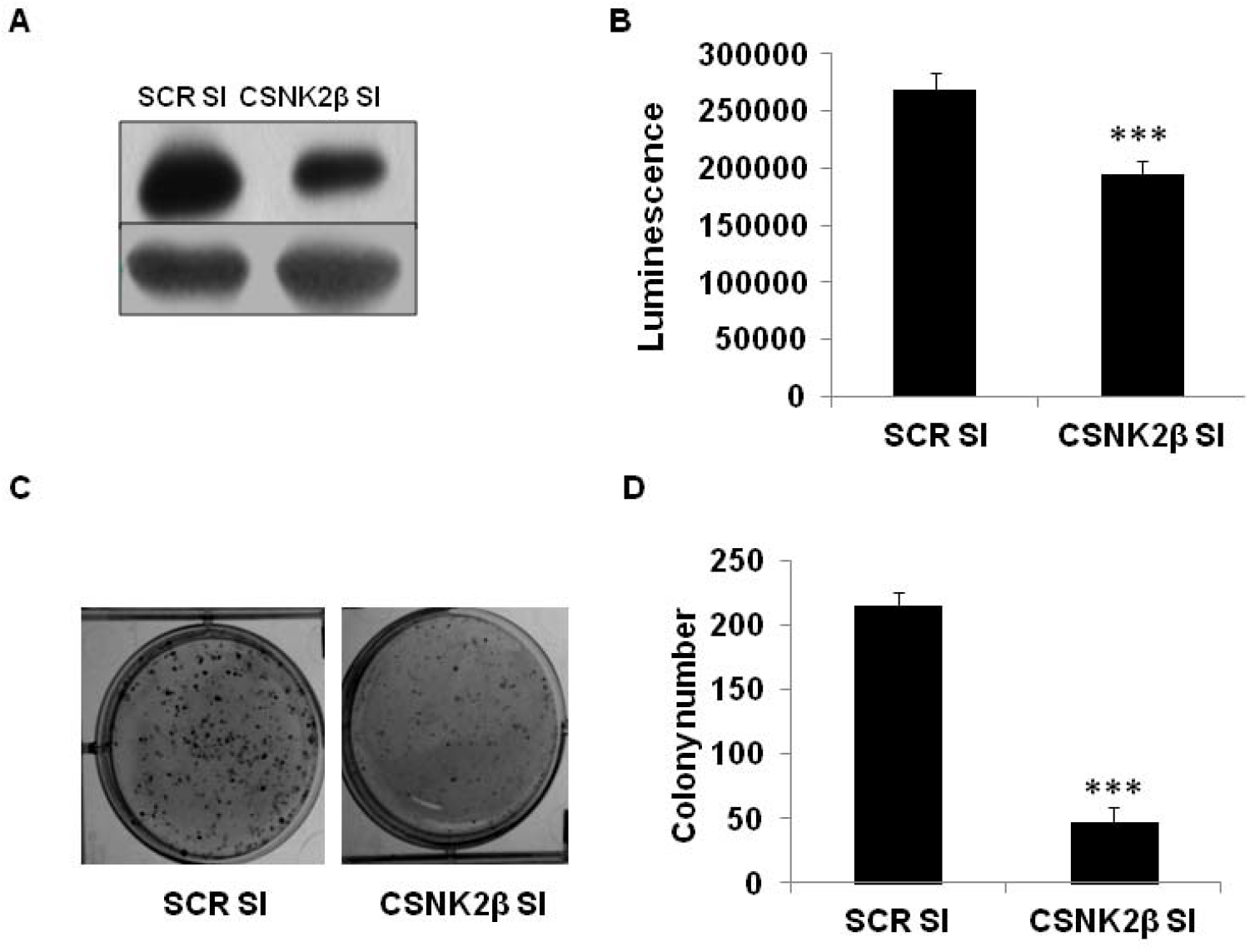
Silencing of CSNK2ß decreases the proliferation and colony formation of MDA-MB-231 cells. (A) Silencing of CSNK2ß gene with siRNA decreased the protein expression of CSNK2ß gene in MDA-MB-231 cells. 2 □ 10^5^ cells were plated in 6 well plates. On the next day, the cells were treated with Scramble and CSNK2ß siRNA and incubated for 72 hours. CSNK2ß expression in the samples was checked by western blotting. (B) Knockdown of CSNK2ß decreases the cell proliferation of MDA-MB-231 cells. 1000 cells were plated in 96 well cell culture plate and reverse transfection was done with respective siRNA and incubated for 96 hours. The cell viability was estimated by CTG method. (C) Knockdown of CSNK2ß decreases the colony formation potential of MDA-MB-231 cells.1000 cells were taken from previously transfected cells with Scramble and CSNK2ß siRNA for 48 h, and seeded in 6 well plates and allowed to grow for 2 weeks. The colonies were fixed with the formaldehyde and stained with crystal violet. The image was taken using gel doc (Biorad) and the colonies were counted using Image J software. (D) Densitometric representation of the number of colonies in each sample. Data are represented as mean ± SD from triplicate samples. ***p < 0.001. SI denotes siRNA in all figures.

### Silencing of CSNK2ß inhibits the colony formation of MDA-MB-231 cells

We also performed the clonogenic assay which associates with the tumor formation *in vivo* [29]. Cells were transfected with Scramble and CSNK2β siRNA in 6 well plates and incubated for 48 hours. After 48 hours of incubation, cells were trypsinized and collected. 1000 cells per well were seeded and allowed to grow until individual colonies were visible (approximately 3 weeks) The colonies were fixed with formaldehyde, stained using crystal violet and counted using Image J software. Silencing of CSNK2β significantly reduced the number of colonies as compared to Scramble siRNA treated samples (p< 0.01) (Figure 2 C, D). This result further confirmed the pivotal role of CSNK2ß in the proliferation of MDA-MB-231 cells.

### Silencing of CSNK2ß attenuates the migratory behavior of MDA-MB-231 cells

Phenotypic assessment of migratory and invasive potential of cancer cells is primary objective before elucidation of underlying mechanism and development of novel strategies for diagnosis; prognosis and therapeutic intervention. Cell motility was determined using classical wound healing assay in which a scratch was made in a monolayer of MDA-MB-231 cells after 48 hours of transfection with scramble and CSNK2β siRNAs. Migratory capacity was observed in each well at different time points (0, 6 and 12h). After 12 hours of wound generation, the area of a wound was significantly wide in CSNK2β transfected wells as compared with scramble suggesting the role of CSNK2ß gene in the migration of MDA-MB-231 cells *in vitro*. (p < 0.05)(Figure 3 A, B).

**Figure 3.**
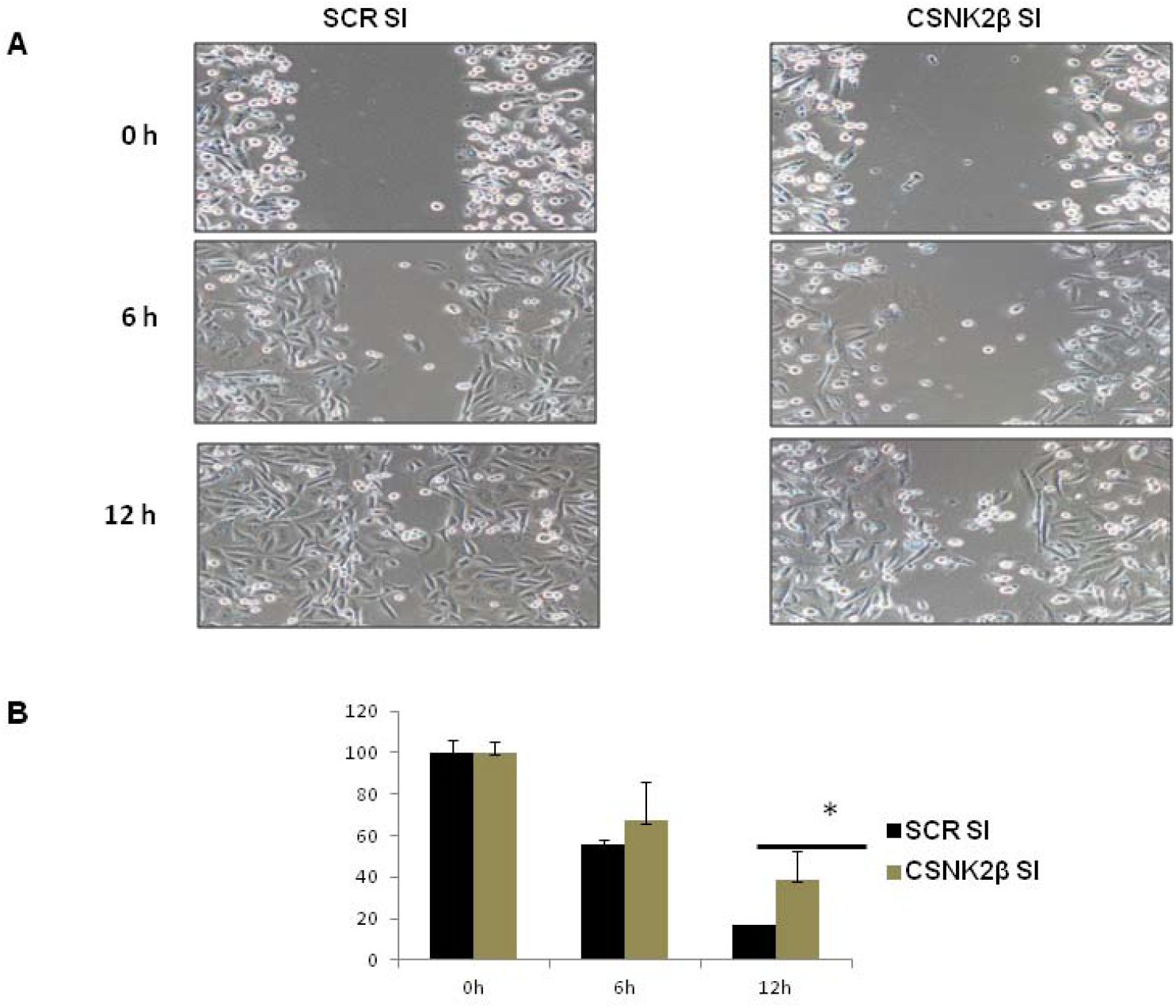
Study of cell migration by wound healing assay. (A) Representative image of wound healing assay done in CSNK2β siRNA treated cells. A wound healing assay was performed to evaluate the motility of cells after silencing CSNK2β for 60 hours. After transfection, a scratch was made on cells monolayer and was monitored with microscopy every 6 hours (0, 6,12h). (B) Densitometric representation of the percentage wound area. The normalized wound area was calculated using Image J software. *p < 0.05.

### Silencing of CSNK2ß induces G2/M arrest in cell cycle analysis of MDA-MB-231 cells

Cell cycle analysis was carried out after knockdown of the cells with Scramble and CSNK2β siRNAs for 48 hours. After 48 hours of post-transfection, the percentage of the cells in the different phase of the cell cycle was analyzed by flow cytometry. The percentage of cells in G2/M phase was significantly increased in CSNK2β transfected samples (22.806%) than in scrambled siRNA transfected samples (13.266%)(p < 0.05). At the same time, the percentage of cells in G0/G1 and S phase was reduced (Figure 4 A, B). These results suggest that knockdown of cells with CSNK2β inhibits the progression of cells via blocking the cell cycle at G2/M phase.

**Figure 4.**
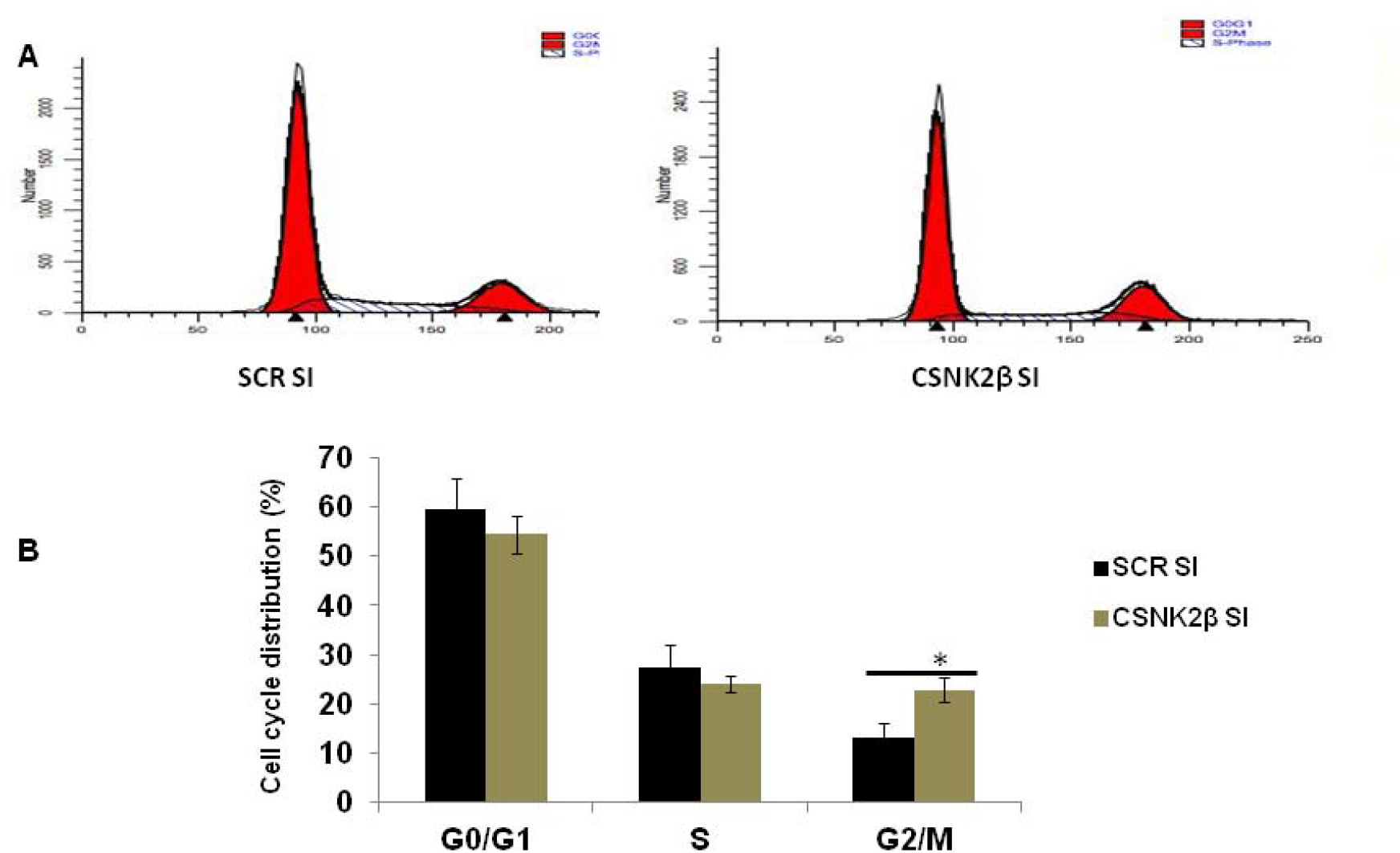
Silencing of CSNK2ß leads to the G2/M arrest. (A) Cell cycle distribution of MDA-MB-231 cells treated with Scramble and CSNK2ß siRNA was evaluated by flow cytometry 48 hours post-transfection, (B) Statistical representation of the percentage of cells in the different phase of cell cycle. Results are representative of three independent experiments represented as a mean ± standard deviation. *p < 0.05.

### Silencing of CSNK2ß induces chromatin condensation indicating apoptotic morphology

After 72 hours post-transfection with CSNK2β siRNA remarkable changes such as chromatin condensation and nuclear fragmentation was seen in the test samples compared to Scramble using Hoechst 33258 staining (Figure 5A). This morphological examination interprets that CSNK2β has a profound role in cell survival and silencing it drives the cells towards apoptosis.

**Figure 5.**
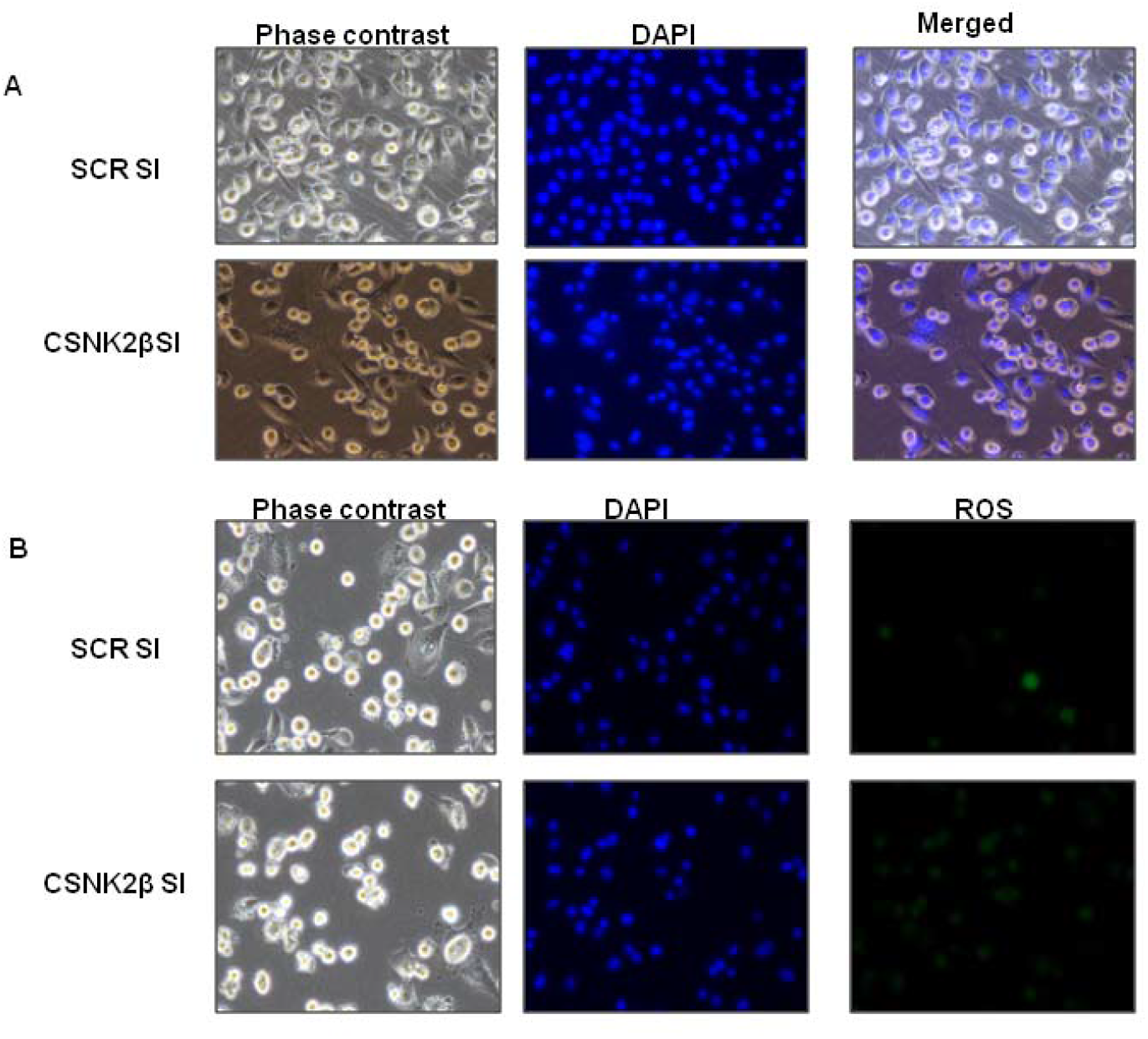
Microscopic evaluation of cell morphology, apoptosis, and intracellular ROS production. (A) MDA-MB-231 cells were observed by fluorescent microscopy after the cells were stained with Hoechst 33342. Cells were transfected with Scramble and CSNK2β siRNA for 72 hours and then stained with Hoechst 33342. (B) For qualitative measurement of intracellular ROS production, the cells were stained with CM-H2DCFDA for 30 minutes in dark and proceeded for Hoechst 33342 staining.

### Silencing of CSNK2ß increases the intracellular ROS production thereby enhancing cell death

ROS possess double edge sword property both as oncogenic, maintaining sustained and increased proliferation of cancer cells as well as tumor suppressor leading to cell death when the aberrant increase in intracellular ROS arises due to any kind of stress [30]. The fine distinction between the ROS activity and the fate of the cells depend on the balance between pro-oxidant and antioxidant or redox status of the cells. It is a good idea to find oxidative stress modulators as an anti-cancer strategy [31]. In our experiment, we found the increased production of ROS in CSNK2β siRNA transfected cells compared to scramble siRNA transfected cells (Figure 5B). This data suggested the CSNK2β modulates the intracellular ROS production in MDA-MB-231 cells.

### Silencing of CSNK2ß induces the cell death of MDA-MB-231 cells by both apoptosis and autophagy mechanisms

To unravel the cell death mechanism induced after the knockdown of CSNK2β, western blot analysis was performed to determine the level of proteins related to apoptosis after 72 hours of post-transfection. We found that there was increased expression of BAX (pro-apoptotic), decreased the level of Bcl-xL (antiapoptotic), activation of procaspase 3 with the increased expression of cleaved caspase 3. (Figure 6 A, B, C). Altogether these results showed that silencing of CSNK2β drives cell death via apoptosis. Moreover, we also analyzed the expression of autophagy markers such as Beclin and LC3. They were both upregulated in CSNK2β siRNA transfected samples than Scramble siRNA. These findings showed that targeting CSNK2ß in MDA-MB-231 triggers the cells towards both apoptosis and autophagy cell death pathway (Figure 6 D, E).

**Figure 6.**
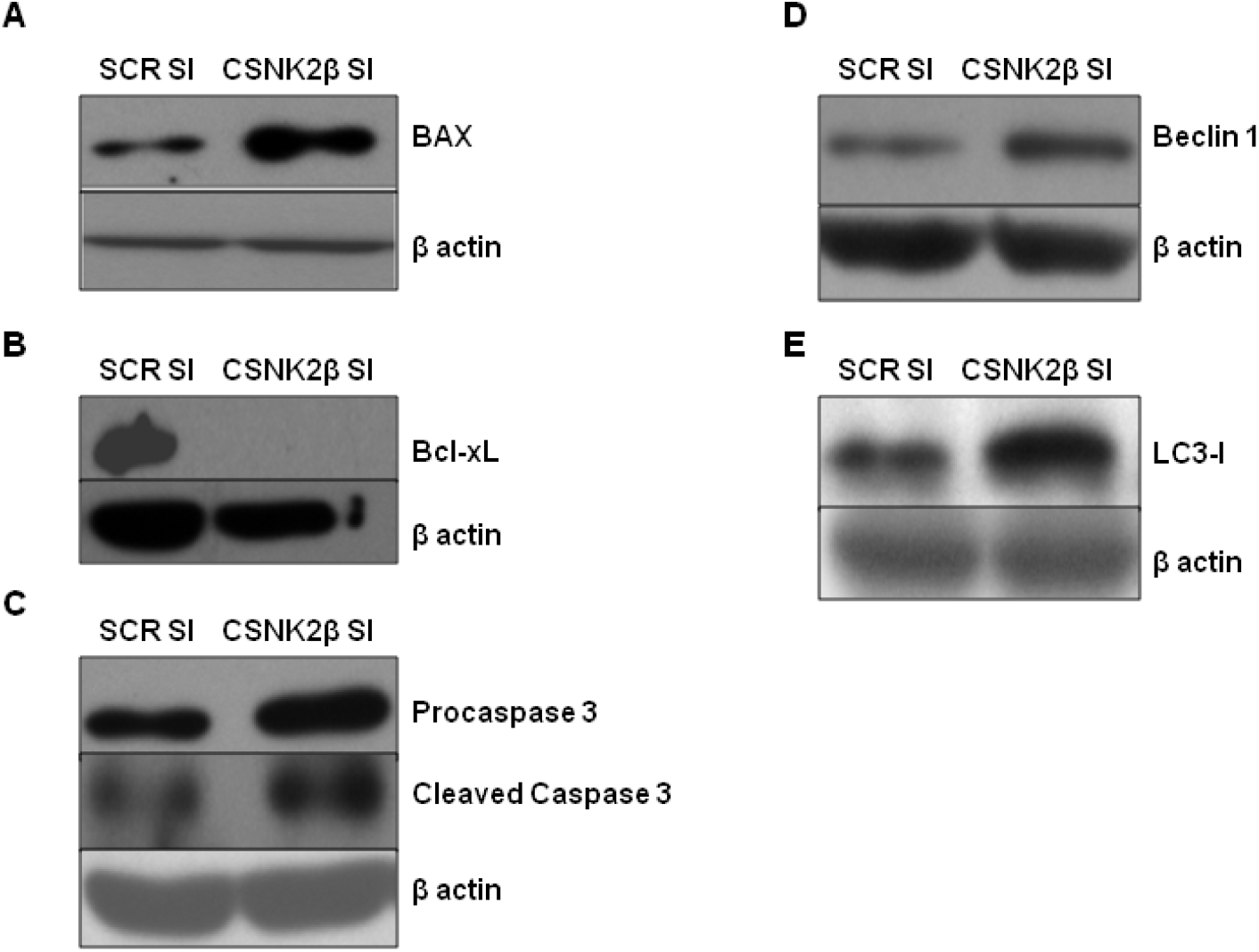
Mechanistic study of the anti-cancer effect of CSNK2ß silencing in MDA-MB-231 cell. (A) MDA-MB-231cells were cultured in 6 well plates (2 □ 10 ^5^ cells/well) and on the next day transfected with Scramble and CSNK2B siRNA. The cells were incubated for 72 hours and RNA was isolated and the level of different genes present in the sample was estimated by using real time PCR. (B-H) The expression level of different proteins related to cell proliferation, survival, and tumor promotion was estimated by western blotting after the cells were transfected with Scramble and CSNK2ß siRNA for 72 hours.

### CSNK2ß regulates the expression of multiple oncogenic signaling pathways

To uncover the gross activity of CSNK2ß in MDA-MB-231 cells, we examined the expression of mRNA and proteins related to cell proliferation, survival, and cell cycle by real time PCR and western blotting. We performed the mRNA expression analysis of a number of genes in 72 hours post transfected samples and found that the mRNA level of p21, a cell cycle inhibitor was increased while the expression of survivin (metastatic marker), Nanog (transcription factor for stemness) and cyclin B1 was decreased following the knockdown of CSNK2β (Figure 7 A). These results suggest the role of CSNK2ß in the cell cycle, cancer stem cell proliferation, and metastasis.

**Figure 7.**
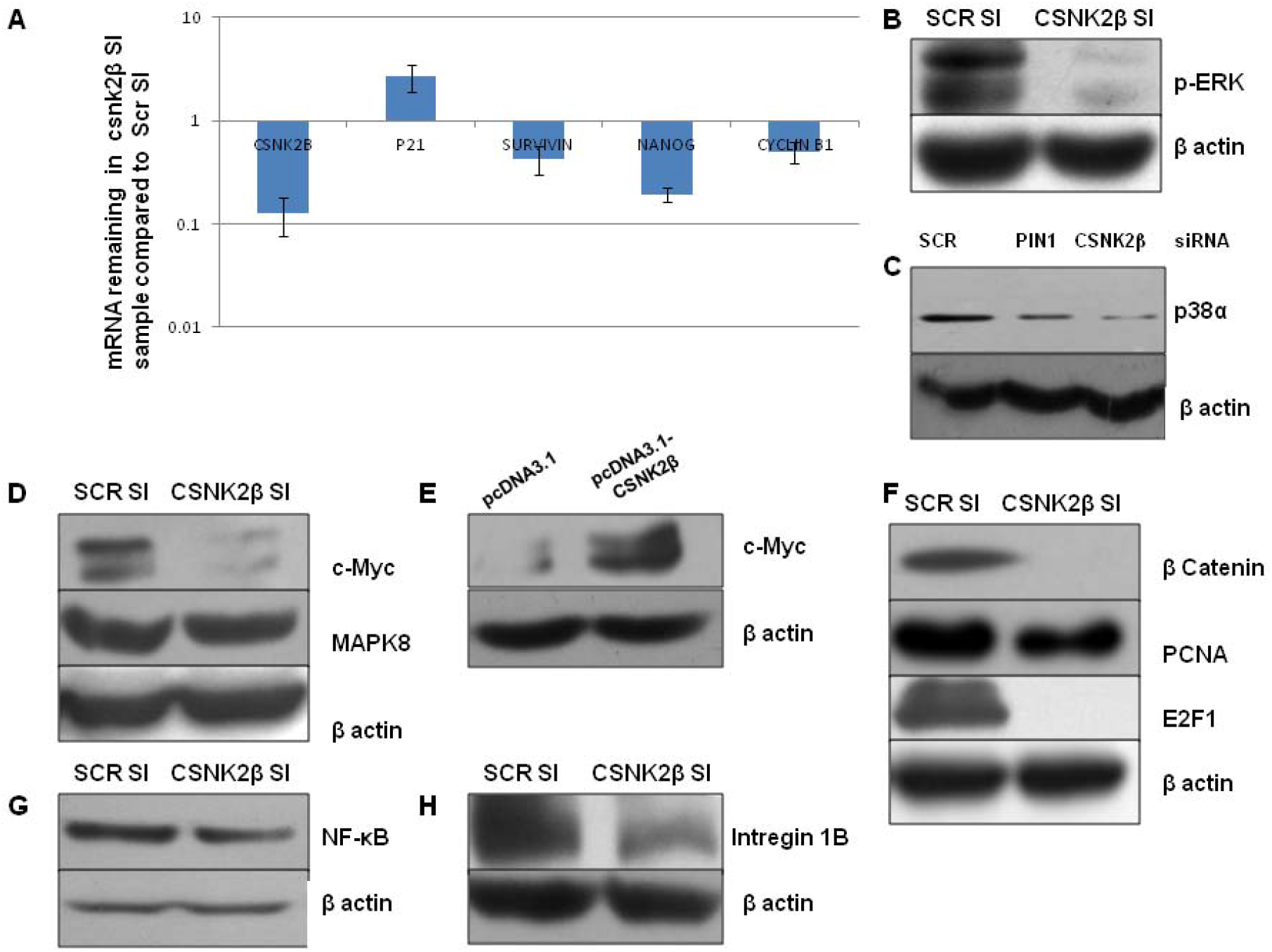
Cell death mechanistic study of CSNK2β silencing against MDA-MB-231 cells. (A-E) Immunoblotting of the Scramble and CSNK2B siRNA transfected samples 72 hours post-transfection was done using apoptosis and autophagy-related antibodies.

To further elucidate the signaling pathways affected by the knockdown of CSNK2β, we studied the expression of proteins responsible for cell proliferation such as p-ERK, p38-α, c-Myc, and MAPK8. CSNK2β downregulation by siRNA transfection inhibits p-ERK, p38-α, c-Myc, and MAPK8 protein expression as compared to Scramble siRNA transfected MDA-MB-231 cells (Figure 7 B-D). Simultaneously Overexpression of CSNK2β increased the expression of c-Myc significantly (Fig 7 E). Hence, our results showed that MAPK pathway is regulated by CSNK2β. Targeting MAPK pathway is prerequisite for the treatment of triple-negative breast cancer (TNBC) [32]. Thus CSNK2β can be the target of choice against TNBC. Further knockdown of CSNK2β also inhibited the expression of β catenin (Wnt signaling pathway), PCNA (DNA replication and repair) and E2F1 (transcription factor) (Figure 7F). Silencing of CSNK2β abrogated the expression of NF-ĸB which is constitutively expressed in breast tumors (Figure 7G) [33]. We also studied the expression of integrin 1 beta, a heterodimeric receptor that senses both external and internal cues and has an important role in tumor biology. CSNK2β siRNA inhibits the expression of Integrin 1β thus decreasing proliferation and invasion (Figure7 H).

### Overexpression and Knockdown of CSNK2β affect the expression of PIN1 and PTOV1

From our previous study, we found that there was the functional relationship between PIN1, PTOV1, and CSNK2β. We have shown that all the three genes share the functional relationship through c-JUN expression that mediated the cancer progression in prostate cancer (PC3 cells). We proposed that these genes fall in the same pathway and mediates their function either by direct interaction with PIN1 or indirectly by promoting the oncogenesis of PIN1 [34]. So we were interested to find the relationship between Pin1–CSNK2β and PTOV1-CSNK2ß in MDA-MB-231 cells. The level of PIN1 and PTOV1 was downregulated with CSNK2β siRNA transfected cells(Figure 8 A,C). Overexpression and knockdown of CSNK2β by pcDNA3.1-CSNK2ß and CSNK2ß siRNA was performed in MDA-MB-231 cells for 72 hours and the expression of PIN1 and c-JUN were measured. Both PIN1 and c-JUN were upregulated with the Overexpression of CSNK2β and decreased with the silencing of CSNK2β. (Figure 7 A, B). Also, we transfected the same clone in HEK239 cells and determine the expression of PTOV1. CSNK2β induces the expression of PTOV1 in HEK239 cells. We also transfected the cells with PIN1, PTOV1, and CSNK2β siRNA and determined the expression of CSNK2β. The level of CSNK2β was not affected in PIN1 and PTOV1 siRNA treated cells which suggested that CSNK2β might be acting upstream of PIN1 and PTOV1.

**Figure 8.**
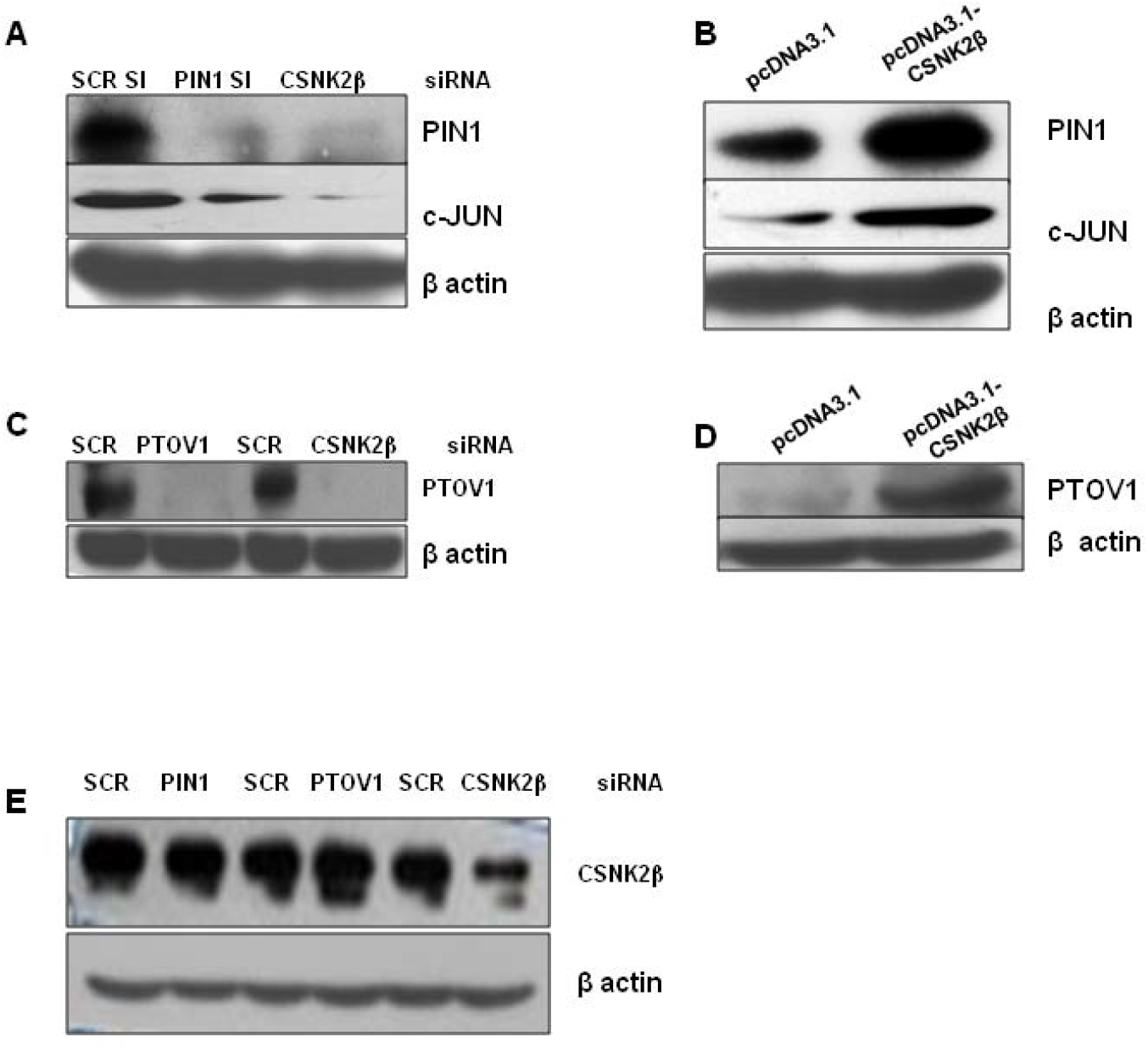
Functional validation of the relationship between CSNK2β, PIN1and PTOV1 in MDA-MB-231 cells. (A-E) Western blot analysis of PIN1, PTOV1, and CSNK2β and c-JUN expression in MDA-MB-231 cells using either PIN1, PTOV1, and CSNK2β siRNA or pcDNA3.1-CSNK2ß.

## Discussion

Breast cancer is the most common cancer occurring in women of both developed as well as developing countries. It was estimated that 1.7 million new cases were diagnosed in 2012 which was 12% of all new cases and 25% of all cancers in women (https://www.wcrf.org/int/cancer-facts-figures/data-specific-cancers/breast-cancerstatistics). Mortality due to breast cancer has been decreasing due to early diagnosis, improved adjuvant therapy, low usage of hormone replacement therapy [35, 36]. Despite all these advancements and outcomes, triple negative breast cancer still remains a major challenge. Statistics show that TNBC represents 10-15% of all breast cancer and has a poor prognosis compared to other subtypes of cancer (ER, PR, and HER2/Neu) [37]. Thus it is indispensable to explore the potential candidate target to reduce the rate of metastases, morbidity, and mortality due to breast cancer and especially TNBC. In our study, we chose to elucidate the role of CSNK2ß in breast cancer (MDA-MB-231, a TNBC cell line) in *in-vitro*.

We first detected the role of CSNK2ß in cell viability and proliferation by CTG and colony formation assay. The results showed that knockdown of CSNK2β significantly decreased the cell viability and colony growth of MDA-MB-231 cells. Further study by wound healing showed that CSNK2β siRNA inhibited the proliferation and invasion of the cells. Silencing of CSNK2β reduces the cell proliferation via arresting the cells at G2/M phase. These all results clarify that CSNK2β has a prominent role in cell proliferation, migration and cell cycle. There is still the controversy about the ROS production and the cell viability and its use as a strategy for anti-cancer therapy. Controlled production of ROS is beneficial for proliferation and viability of cancer cells but its elevated level may be genotoxic and cause apoptosis of cancer cells [30, 31]. Microscopic study of CSNK2β silenced MDA-MB-231 cells showed the increase in ROS production and nuclear condensation. Both the features support that the cells were in the process of apoptosis after the silencing with CSNK2β siRNA. Studies on different isoforms of casein kinase and their subunits in eukaryotic biology and cancer have been studied [1–3], but the possible mechanistic study of CSNK2ß in cancer was unclear. Although CSNK2ß is a regulatory subunit of CSNK, it is highly conserved protein and possesses independent function. Thus we anticipate that finding the new molecules within the CSNK2β regulatory network will tease out its role in tumorigenesis. To get the further insight into the biological significance of this protein we performed the expression study of the key molecules related to cell proliferation, survival, cell cycle, apoptosis and autophagy-related genes by western blotting and real time PCR.

Although the improved chemotherapy and use of adjuvants have decreased death rates due to breast cancer, understanding the response to treatment and apoptosis is still a major concern [38]. Here we have shown the induction of BAX (proapoptotic), suppression of Bcl-xL(antiapoptotic), activation of procaspase 3 with an elevated level of cleaved caspase 3 in CSNK2β siRNA transfected sample suggesting the cell death via apoptosis. Paradoxically autophagy is utilized as a survival mechanism by tumor; it can also promote the caspase-independent form of cell death and can be used as a cancer treatment modality [39, 40]. Moreover, BECN1 and LC3 which are autophagy markers were increased in CSNK2β knockdown samples. Since it follows both apoptosis and autophagy cell death mechanism, CSNK2ß is an interesting molecule for targeted cancer therapy.

Raf/MEK/ERK pathways are activated in many tumors (prostate, breast, leukemia, melanoma, thyroid) which transmit the signals from cell surface receptors to transcription factors and can be exploited for therapeutic intervention [41,42]. In the present study, we found that CSNK2β regulates the expression of P-ERK, p38-α, c-MYC, MAPK8 proteins which are connected to MAPK pathway in MDA-MB-231 cells. These data suggested that targeting CSNK2β might be a potential strategy to improve clinical outcomes in future. Numbers of studies have been carried out on the dysregulation of Wnt/β-catenin in human breast cancer and is good clinical and pathological marker with the poor survival outcome [43, 44]. Consistent with these reports, our data also showed that knockdown of CSNK2β down-regulates the expression β-.catenin. PCNA is earlier reported as a reliable marker to access the growth and predicting the prognosis in breast cancer [45]. Thus we investigated the effect of silencing of CSNK2β on PCNA and its downregulation supported the previous findings. A recent study on the interbreeding of MMTV-PyMT mice with E2F1, E2F2, or E2F3 knockout mice showed that in addition to cell cycle control E2F targets the number of genes related to angiogenesis, extracellular matrix modification, proliferation and survival of tumor cells which was important for metastasis [46]. We also found that CSNK2β reduces the expression of E2F1 which might affect a large fraction of genes in breast cancer. NF-ĸB expression leads to the induction of genes related to apoptosis, cell cycle, cell invasion which contributes to tumorigenesis, chemoresistance and radioresistance [47]. Our result showed that CSNK2β might regulate the cell proliferation through the NF-ĸB pathway. From wound healing experiment we deduce that CSNK2β has a remarkable role in cell migration. This was further supported by the decrease in integrin 1B, glycoprotein receptors that mediate anchorage and migration of cells via cell matrix and cell-cell interactions [48].

PIN1, a widely established oncogene is overexpressed in the breast cancer. PIN1 increases the expression of cyclin D1expression by different pathways activating the c-Jun, β catenin/T cell transcription factor (TCF) and NF κB transcription factors [49,50]. In continuation of our previous study [34], we have shown that overexpression of CSNK2β increases the expression of PIN1 while silencing it decreases its expression. Although the study on PIN1 and CSNK2α is reported previously our results provided a new avenue for further exploration. Also, we have shown that Overexpression of PIN1 and CSNK2β elevated the level of C-Jun and Knockdown by siRNA reduced the expression. There are several reports of PTOV1 and human breast cancer. Overexpression of PTOV1 promotes tumor progression and is a predictor of poor prognosis in breast cancer [51]. Also, PTOV1 promotes the tumorigenicity by activating Wnt/β-catenin pathway in breast cancer [52]. In our study, we have shown an initial report that overexpression and knockdown of CSNK2β both increased and decreased the expression of PTOV1respectively.Our previous study on co-immunoprecipitation showed that PIN1 interacts with PTOV1. These results support our previous finding that PIN1, PTOV1 and CSNK2β fall in the same pathway and either directly interact or cooperatively promote the oncogenesis [34].

This conjecture provides a new scope for further experiments and validations to establish the relationship between these molecules and decipher their role in cell proliferation, survival, migration, and cell death in breast cancer as well as other cancer models.

## Conclusion

In summary, our findings from our laboratory have shown that silencing of CSNK2β with siRNA significantly reduced the cell viability, colony formation potential, cell migration and proliferation of MDA-MB-231 cells. CSNK2β regulates the expression of proteins related to MAPK, JNK, Wnt, apoptosis, and autophagy signaling pathways. Silencing of CSNK2β arrests the cells in G2/M phase, causing the decrease in cell proliferation. Our study has provided the first evidence that silencing CSNK2β decreased the expression of PIN1 and PTOV1 oncogenes previously studied in breast cancer. Taken together, our study not only showed the importance of CSNK2β as a novel target for breast cancer therapy but also suggest therapeutic implications of targeting CSNK2β with siRNA or other approaches.

## Declarations

### Acknowledgements

We would like to thank Professor Jukka Westermarck for guiding our research work on CSNK2β

### Funding

SAU-START-UP-GRANT-2013, South Asian University, New Delhi, India. Funding sources has no role in the design of the study, analysis of data, and writing the manuscript.

### Availability of data and materials

Data supporting these results are available from the authors upon request.

### Authors’ contribution

SKLK performed most of the work reported, including writing the manuscript, performing CTG, CFA, qRT-PCR, WB, Wound Healing, Cell Cycle analysis etc. BAL designed clones and confirmed their expression. FA performed ROS and Hoechst staining, NS helped in performing WB and YRP designed the study, provided overall direction, oversight and writing manuscript of the project. All authors read and approved the final manuscript.

### Ethics approval

Not applicable

### Consent for publication

This manuscript does not contain any individual person’s data in any form (including individual details, images or videos).

### Competing interests

The authors declare that they have no competing interests.

